# Constraint-based modeling predicts metabolic signatures of low- and high-grade serous ovarian cancer

**DOI:** 10.1101/2023.03.09.531870

**Authors:** Kate E. Meeson, Jean-Marc Schwartz

## Abstract

Ovarian cancer is an aggressive, heterogeneous disease, burdened with late diagnosis and resistance to chemotherapy. Clinical features of ovarian cancer could be explained by investigating its metabolism, and how the regulation of specific pathways link to individual phenotypes. Ovarian cancer is of particular interest for metabolic research due to its heterogeneous nature, with five distinct subtypes having been identified, each of which may display a unique metabolic signature. To elucidate metabolic differences, constraint-based modeling (CBM) represents a powerful technology, inviting the integration of ‘omics’ data, such as transcriptomics. However, many CBM methods have not prioritised accurate growth rate predictions, and there are very few ovarian cancer genome-scale studies, thus highlighting a niche in disease research. Here, a novel method for constraint-based modeling has been developed, employing the genome-scale model Human1 and flux balance analysis (FBA), enabling the integration of *in vitro* growth rates, transcriptomics data and media conditions to predict the metabolic behaviour of cells. Using low- and high-grade ovarian cancer as a case study, subtype-specific metabolic differences have been predicted, which have been supported with CRISPR-Cas9 data and an extensive literature review. Metabolic drivers of aggressive phenotypes, as well as pathways responsible for increased proliferation and chemoresistance in low-grade cell lines have been suggested. Experimental gene dependency data has been used to validate fatty acid biosynthesis and the pentose phosphate pathway as essential for low-grade cellular growth, highlighting potential vulnerabilities for this ovarian cancer subtype.

## Introduction

Ovarian cancer (OC) is the seventh most common cancer in women, accompanied by a staggering 1 in 75 lifetime risk (Momenimovahed et al., 2019), (Reid et al., 2017). It is typically diagnosed at late-stage, where less than 1 in 3 women will survive 5-years, thus ovarian cancer signifies a major clinical burden (Reid et al., 2017). Although it may also develop from sex-cord stromal or germ cells, over 90% of ovarian cancers are epithelial in origin (Reid et al., 2017). Furthermore, ovarian cancer itself is an extremely heterogeneous disease, which may be split into five main subtypes: high-grade serous (HGSOC), endometrioid, clear cell, mucinous and low-grade serous (LGSOC) (Jayson et al., 2014). High-and low-grade serous ovarian cancer represent subtypes with contrasting phenotypes and genomic profiles. HGSOC – the more aggressive subtype – makes up around 70% of cases, with ubiquitous TP53 mutations, homologous recombination deficiency in 50% of cases and BRCA1/2 inactivation (Reid et al., 2017), (Shih & Kurman, 2004), (Jayson et al., 2014). On the other hand, LGSOC constitutes less than 5% of ovarian tumours, and is defined by BRAF, KRAS and PTEN mutations (Reid et al., 2017) (Jayson et al., 2014).

Standard, first-line treatment for ovarian cancer involves cytoreductive, debulking surgery, accompanied by platinum-based chemotherapy, such as cisplatin or paclitaxel (Cortez et al., 2018). The heterogeneity of ovarian cancer complicates treatment strategies, with HGSOC demonstrating a more positive initial response, however too often patients will relapse (Jayson et al., 2014). Although distinguishable by their histology and pathogenesis (Vang et al., 2009), if metabolic characterisation was also considered, this could permit a more detailed prognosis, given metabolism is directly linked to therapeutic response and the mechanisms of cancer progression.

Metabolic features of high-versus low-grade serous ovarian cancer have been reviewed in recent years, revealing distinguishing features of carbon and energy metabolism, as well as fatty acid metabolism. For example, glycolysis has always been a central focus regarding cancer metabolism, and although the Warburg effect (preference for aerobic glycolysis even in the presence of oxygen) has been recognised as a Hallmark of cancer, it has been suggested that in some subtypes, oxidative phosphorylation is the preferred pathway (Warburg et al., 1927), (Nantasupha et al., 2021). Furthermore, the rate of flux through glycolysis may differ between high-and low-grade serous ovarian cancer, and high-grade tumours might upregulate oxidative phosphorylation to a greater extent (Nantasupha et al., 2021). Aside from carbon metabolism, numerous enzymes involved in fatty acid metabolism have been identified as key drivers, for example the overexpression of fatty acid synthase (FASN) has been associated with poor prognosis in OC, amongst other enzymes such as ATP-citrate lyase and acetyl-CoA carboxylase (Ji et al., 2020).

Genome-scale models (GEMs) are efficient for transfer of in-vitro results (such as gene and protein expression data) onto an in-silico platform. In principle, GEMs can be thought of as a multi-dimensional network, containing all known metabolic reactions and associated enzymes for a particular organism, serving as platforms onto which transcriptomics data can be integrated to create a context-specific, personalised GEM. The process of producing these enzyme-constrained GEMs (ecGEMs) is called constraint-based modeling (CBM), and such ecGEMs can be manipulated via genetic engineering simulations, such as gene overexpression or deletion, reaction inhibition, essentiality predictions and drug simulations (Ebrahim et al., 2013). Thereby, ecGEMs enable us to make predictions to be tested with experimental work, which could save time and resources, potentially aiding the decision-making and hypothesis-generating process. In an ideal situation, a human GEM could be constrained with RNAseq data gained from a patient’s tumour, and used to predict therapeutic responses and disease progression, directing treatment decisions and helping generate more accurate prognoses. Genome-scale modelling poses a higher predictive potential than the direct measurement of metabolic fluxes, as *in vitro* experiments are limited to measuring only a few dozens of reactions in central metabolism (Yizhak et al., 2015). Thereby the hypothesis-generating capability of wet lab methods may be complemented by modelling approaches, providing a worthwhile link between abundant omics data and reaction rate predictions.

Over the past couple of decades, studies into human metabolism have been greatly advanced by GEMs: namely, the Recon series (Recon1, 2 and 3D) (Duarte et al., 2007), (Thiele et al., 2013), (Brunk et al., 2018) and the Human Metabolic Reaction series (HMR1 and 2) (Mardinoglu et al., 2013), ((Mardinoglu et al., 2014). In addition to these fundamental models there have been whole-body GEMs, for example two complete reconstructions, containing organ- and sex-specific information (Thiele et al., 2020), from which individual organs could be extracted. Utilising HMR2 as its framework, one project constrained six patient-specific GEMs, which enabled the discovery of anticancer drugs (Agren et al., 2014). One of the most recent human GEMs to be developed was the Human1 model, which is an updated and curated version of the Recon and HMR series, demonstrating optimisation such as the removal of duplicated reactions and metabolites, correction of model inconsistencies, and the standardisation of identifiers (Robinson et al., 2020). There are many existing integration algorithms to constrain these GEMs with omics data. Some of these utilise discrete methods, switching reactions on or off according to expression thresholds, for example iMAT (Zur et al., 2010) and GIMME (Becker & Palsson, 2008). On the other hand, some algorithms use continuous bounds, where expression values are applied directly, such as E-Flux (Colijn et al., 2009). However, genome-scale modeling of ovarian cancer remains limited; one well-cited study reporting the constraining of Recon1 to decipher resistance mechanisms in ovarian cancer did not provide *in vitro* validation of growth rates or metabolic predictions, nor was an original GEM or omics integration algorithm developed (Motamedian et al., 2015).

Here, a novel omics integration algorithm is presented, which accurately tailors the Human1 genome-scale model towards *in vitro* growth measurements, using experimental growth rates as internal thresholds. The proposed algorithm involves the use of continuous, relative reaction bounds – incorporating normalised transcriptomics data directly. Furthermore, the low- and high-grade ovarian cancer-specific models presented here provide a framework for genetic engineering experiments, which could generate interesting hypotheses for future work, as well as thought-provoking predictions on the difference between low- and high-grade ovarian cancer metabolism.

## Methods

### CCLE transcriptomics data

Transcriptomics data were obtained from the Broad DepMap Portal, under the DepMap Public 22Q2, titled ‘CCLE_expression_full.csv’ (https://depmap.org/portal/download/all/). Out of tens of primary diseases, this dataset includes 65 ovarian cell lines and 53,949 genes (accessed September 2022). This dataset contains RNAseq TPM gene expression data for all genes using (RNA-Seq by Expectation-Maximization) RSEM, which has been Log_2_ transformed, using a pseudo-count of 1, in order to avoid negative values. Further normalisation and processing details were described alongside dataset release (Ghandi et al., 2019). As acknowledged by a similar study utilising CCLE data for genome-scale modeling (Vieira et al., 2022), the relationship between RNA and proteins is complex and not completely understood in the context of its constraint-based modeling implications, however there is a moderate positive correlation between gene and protein expression (r = 0.48) (Nusinow et al., 2020). Therefore, in this study, gene expression was used as an alternative for protein expression.

### Cell line annotations and growth conditions

Before models could be built, it was necessary to annotate the CCLE cell lines with their subtypes, so sample groups could be chosen. Due to the fact there is contention in literature as to the ‘true’ subtype of some ovarian cell lines, we have chosen to use recent NMF clustering to direct our subtyping (Barnes et al., 2021), because this paper combined both an extensive literature review and a comprehensive NMF clustering workflow to determine the true subtypes of cell lines, demonstrating high diagnostic accuracy on patient-derived models and validating previous classification analyses. Media conditions have been reported by Barrentina *et al*. (Barretina et al., 2012), and where these weren’t available for specific cell lines, the source of optimal media, as well as the site of origin, has been referenced in Supplementary file 1 (‘lg_hg_media_subtype_site’).

### Human1 genome-scale model and Metabolic Atlas

For this project, constraint-based modelling techniques were applied to the Human1 model which was obtained from the Human-GEM GitHub repository (https://github.com/SysBioChalmers/Human-GEM) (Robinson et al., 2020), (H. Wang et al., 2022). The model version used here is titled ‘Human-GEM-annotated.xml’ and was accessed in September of 2020. This version includes 13,096 reactions, and of these, 61.4% are annotated with gene-reaction rules, informing the enzyme(s) required for catalysis. These gene-reaction rules are where transcriptomics data has been integrated. Of this annotated portion, there are 653 ‘and’ rules, which specify enzyme subunit IDs; there are 3972 ‘or’ rules, which describe isoenzymes; there are 129 ‘andor’ rules which include both isoenzymes and subunits, and the remaining 3282 annotated reactions are ‘one-gene’ rules, which list one single enzyme needed for catalysis. In addition, there are 3628 unique genes in the Human1 model, and of this total, there are only four which are not included in the CCLE transcriptomics dataset (RNF115, RNF128, SLC27A2 and METTL23). The units for the Human1 model are mmol/gDW/hour for all reactions, except for biomass production, which is g/gDW/hour, therefore the inverse of growth rate corresponds to doubling time.

Metabolic Atlas is the accompanying web portal, which was published alongside the Human1 model (Robinson et al., 2020), and has been used to understand and contextualise the cell line-specific results generated from this project.

### Constraint-based modeling and transcriptomics integration workflow

All methods described below, alongside accompanying requirements and dependencies, have been made available on the Github repository ‘ovarian_genome_scale_modeling’ (https://github.com/katemeeson/ovarian_genome_scale_modeling). This method employs FBA, and has been optimised for biomass production, a widely used cellular objective function for the constraint-based modeling of cancer cells (Yizhak et al., 2015). In order to integrate transcriptomics data, MEWpy, COBRApy and complementary original code was used to generate a novel workflow. The preliminary step for data integration was to define media conditions, involving exchange and demand reactions, and then normalised gene expression values were integrated with the bulk of the Human1 model, in a cell line-specific manner. For all simulations, the GLPK solver and Python 3.0 were used (Van Rossum & Drake, 2009).

### Validating model predictions

In order to validate model predictions, an over-representation analysis (ORA) was performed in WebGestalt, using the ‘pathway’ and KEGG functional database and the ‘genome’ reference set (Liao et al., 2019). The input for this search was experimental gene dependency data, which was calculated from CRISPR knockout (KO) screens and describes how essential a gene is for growth. The gene dependency dataset was accessed from the CCLE Broad DepMap Portal, under the DepMap Public 22Q2, and titled ‘CRISPR_gene_dependency.csv’. From this dataset, a subset was extracted containing the same cell lines on which modeling was performed, namely, three high-grade (COV318, CAOV3 and OAW28) and two low-grade cell lines (59M and HEYA8), however the low-grade OV56 cell line was not available. From here, average gene dependencies were calculated for the two low-grade cell lines, and three high-grade, then these mean values were compared. The 200 genes which showed the highest increase in dependency in low-compared to high-grade cell lines were used as input for the WebGestalt ORA, in order to deduce the overarching pathways which low-grade cell lines might be relying on for growth.

Results were also validated by comparing the growth effect of *in silico* KO simulations set up using COBRApy (Ebrahim et al., 2013), with the CRISPR experimental data described above. A ratio of growth after single gene deletion over optimal growth was calculated and recorded for all 3628 genes in the model. From here, genes present in both the CRISPR dataset and model simulations (n = 3417) had their gene dependency score and KO simulation growth ratio correlated. Due to the expected linearity between *in silico* growth ratio and *in vitro* gene dependency, and given the dependency score was originally calculated as a function of growth, Pearson correlation coefficient was used. Following an analysis of all 3417 genes, a subset of genes with predicted high essentiality were correlated, namely those with a growth ratio of less than 1.0.

## Results

### Standardisation of media composition

In order to build this novel transcriptomics integration algorithm, three low-grade (59M, HEYA8, OV56) and three high grade serous ovarian cell lines (CAOV3, COV318, OAW28) OC cell lines were used, as visualised in Figure 1.

**Figure 1.**
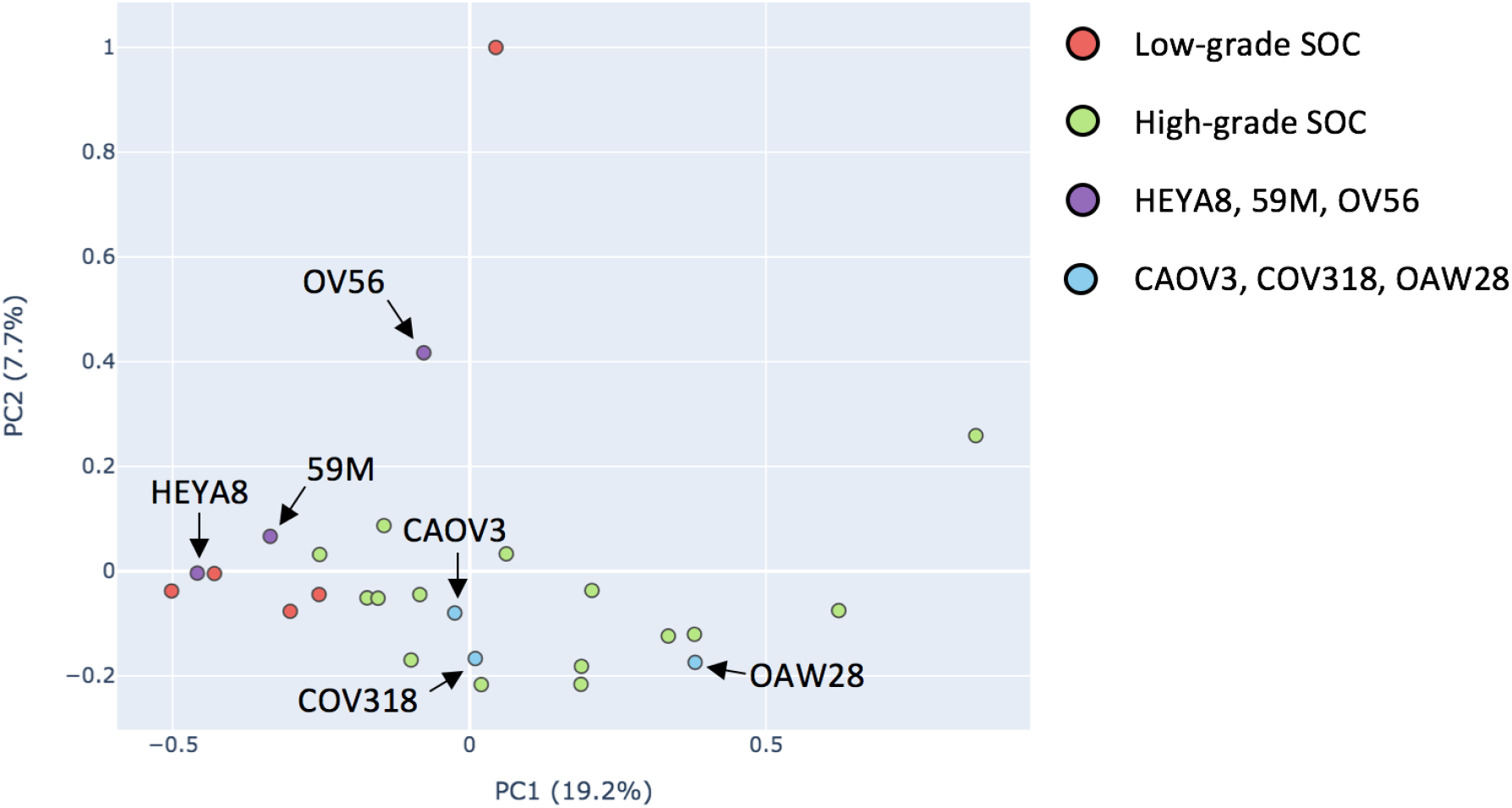
Visualisation of selected cell lines. 2D Principal components analysis visualising the spread of the CCLE RNAseq low- and high-grade serous ovarian cancer (SOC) cell lines, with chosen cell lines highlighted. ‘True’ labels were assigned according to Barnes *et al*., 2021. Green (and highlighted blue) are high-grade, and red (and highlighted purple) are low-grade serous OC. Data from CCLE (accessed September 2022).

In order to direct the selection of cell lines for sample groups, the entire subset of CCLE low- and high-grade serous OC cell lines was visualised using PCA and labelled according to their subtype, media conditions and site of origin (Supplementary Figure 1). The purpose of this visualisation was to deduce whether or not the contrasting media conditions and site of origin directly affected gene expression, and if so, the acceptable variation of media composition and site of origin for the cell lines to be compared in this study. The constrained models had their media completely standardised to DMEM + 10% FBS – the simplest media definition available. CAOV3 had its media systematically changed to assess how this affected the spread of points on PCA (Supplementary Figure 2). These comparisons showed that whilst DMEM remained a main component of the media, metabolic flux results did not change, as shown by the general PCA-visualised spread of the cell lines. In addition, cell lines were chosen from the central regions of either subtype cluster (low- or high-grade) (see Figure 1), to help tailor sample groups to yield subtype-specific constrained GEMs, provided metabolic flux results are in proportion to gene expression data

### Defining media conditions within Human1 model

The full workflow for data integration has been illustrated in Figure 2. The media conditions for each cell line are defined as a preliminary step, and have been described in Supplementary table 1.These were integrated using the MEWpy ‘get_simulator’ function, setting the final media dictionary as the ‘envcond’ argument (Pereira et al., 2021). A limitation of defining media for human GEMs is the inclusion of fetal bovine serum (FBS), which is a complex, difficult to define solution. Therefore, FBS components were estimated by reopening specific reactions according to predicted essentiality (relaxing bounds back to original −1000 or 0 and +1000 mmol/gDW/hour); for example, the exchange reaction ‘MAR09107’, which imports heme into the cytosol, was reopened in the HEYA8 media dictionary. Heme is suggested to be present in FBS preparations at a concentration between 1.1 and 2.0μM (Wagner et al., 2007), which validates the reopening of the heme-importing reaction in media dictionaries. Final copies of media-constrained models were saved and utilised in the transcriptomics integration stages described below.

**Figure 2.**
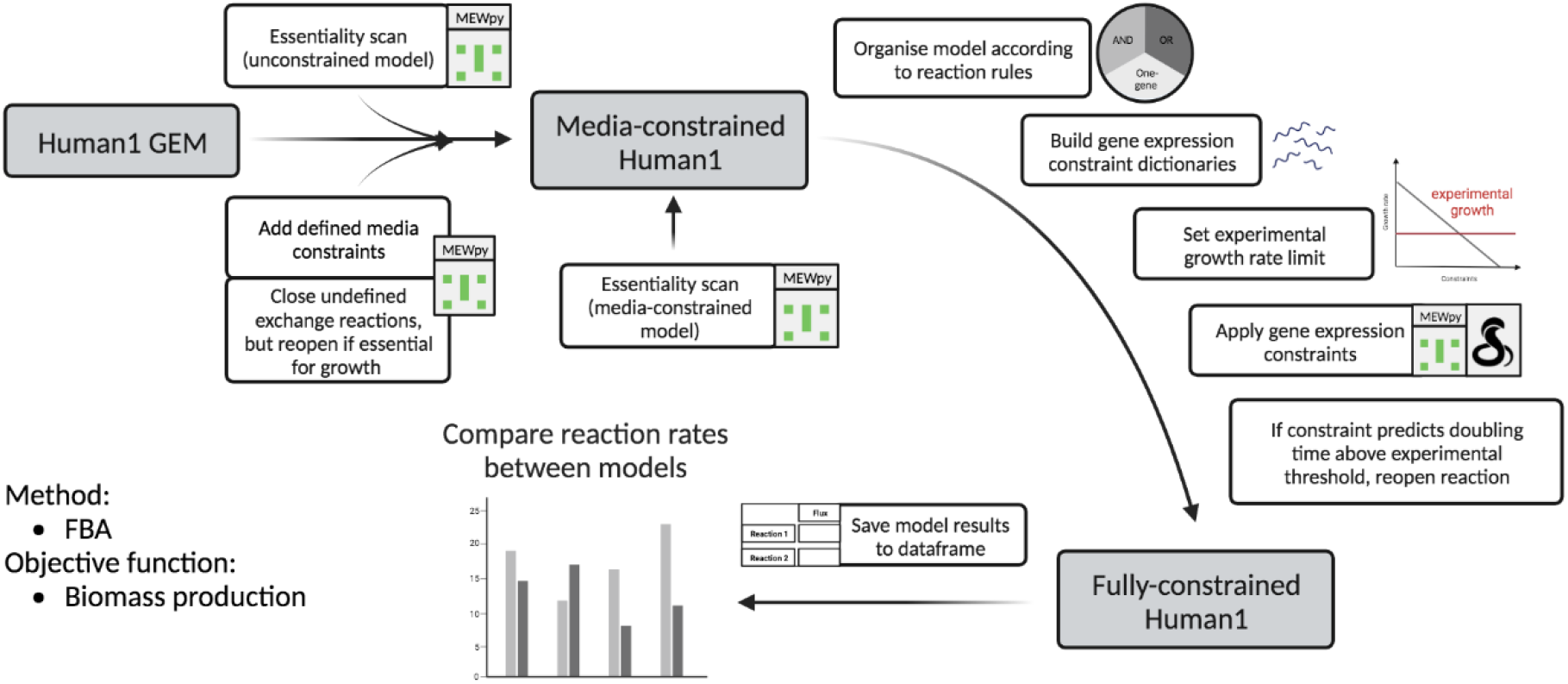
The transcriptomics integration workflow. First, a media dictionary was built, using defined components and closing all other exchange reactions, unless MEWpy predicts positive essentiality. Media constraints were applied to unconstrained Human1 model, using MEWpy. A gene essentiality scan was performed on the media-constrained model. Normalised transcriptomics data was organised into dictionaries, and applied to the media-constrained Human1 model, using MEWpy ‘get_simulator’ function and ‘constraints’ argument. Gene expression constraints were applied according to reaction rules, as described in Table 1. Reactions were reopened if gene expression constraints predicted doubling time over experimental threshold. The method used was FBA, with biomass production as the objective function. Flux results were saved to a dataframe for ease of comparison between cell line-specific GEMs. Graphic created on Biorender.com.

### Integrating transcriptomics data into Human1 model

This novel integration method is based on the incorporation of expression data based on cross-reference with Human1 gene-reaction rules, which define the enzymes and their genes required for reaction catalysis. Every gene reaction rule contained within Human1 has a lower and upper bound, with default excess reaction rate limits of −1000 and +1000, or 0 and 1000 mmol/gDW/hour, based on encoded reaction reversibility. Whether or not the expression data is applied directly, as a sum or as a minimum value depends on the category of reaction rule, as described in Table 1. Reaction rules using the ‘andor’ term have not been included in this integration method, due to the fact they only represent 1.6% of the total annotated portion of the model.

**Table 1.** Method of rule-dependent expression data integration. If the Human1 reaction rule specifies a single gene, then the normalised expression data can be used directly as the reaction bounds. If an ‘or’ term is specified (and no ‘and’ term), then the normalised abundances of either isoenzyme can be summed, as the probability of the reaction being catalysed has been assumed to be directly proportional to the cumulative amount of compatible isoenzymes. If there is an ‘and’ term specified (and no ‘or’ term), then the minimum expression value of all available subunits is taken as the reaction bounds, as this has been assumed to be the rate limiting value. A and B denote reactants; C denotes product; E denotes reaction-specific enzyme; E_1_ and E_2_ denote isoenzymes, and E_a_ and E_b_ denote enzyme subunits. Where expression value ‘or 0’ has been specified for the lower bound in the final column, this refers to a reaction which may occur in the reverse direction, or forward direction only, respectively.

| Human1 reaction rule category | Gene-reaction rule schematic | Method of integrating expression data |
| --- | --- | --- |
| One-gene | $A + B \xrightarrow{E} C$ | Lower bound = -E or 0<br>Upper bound = +E |
| ‘or’<br>(isoenzymes) | $A + B \xrightarrow{E_1 \text{ or } E_2} C$ | Lower bound = -E<sub>1</sub> + -E<sub>2</sub> or 0<br>Upper bound = E<sub>1</sub> + E<sub>2</sub> |
| ‘and’<br>(enzyme subunits) | $A + B \xrightarrow{E_a \text{ and } E_b} C$ | Lower bound = min(-E<sub>a</sub>, -E<sub>b</sub>) or 0<br>Upper bound = min(E<sub>a</sub> + E<sub>b</sub>) |

In parallel to the method described above for defining media conditions, transcriptomics data was integrated with a combination of reaction reopening and closing, based on genes’ inclusion in the CCLE expression dataset and essentiality predictions. It has been reported that gene and protein expression within the CCLE dataset have low correlation (Nusinow et al., 2020), and also that many omics integration algorithms, such as iMAT, do not accurately predict growth rates as they do not account for metabolic deviation or adaptation from a transcriptomics signature (Jerby et al., 2012). Therefore, here, experimental growth rates have been used as a limit for model-predicted growth, and we reopened any reactions which forced growth rates beyond this limit. This means experimental growth thresholds are serving to estimate where gene expression does not accurately represent enzyme abundance and predict experimentally-derived growth rates. A general workflow for this gene expression integration has been illustrated simplistically in Figure 2.

### Using experimental maximum doubling times to tailor model-predicted growth rates

Prior to data integration, the flux through biomass production was approximately 187 g/gDW/hour. Following the standardisation of media composition described above, and integration of these constraints, the doubling time dropped to between 4.5 and 5 hours for every model (equivalent of between 0.2 to 0.22 g/gDW/hour) (Figure 3). Although a lot more realistic than the original, unconstrained solution, these doubling times were still far too high compared to experimental values, and needed supplementing with gene expression constraints to reach a more realistic value. The experimental growth rates, as outlined in Supplementary table 2 (‘Experimental and model-predicted growth rates’), were input as the minimum growth rate threshold which the corresponding cell line-specific model could reach, by writing code which reopened any reactions constraining biomass production under this threshold.

**Figure 3.**
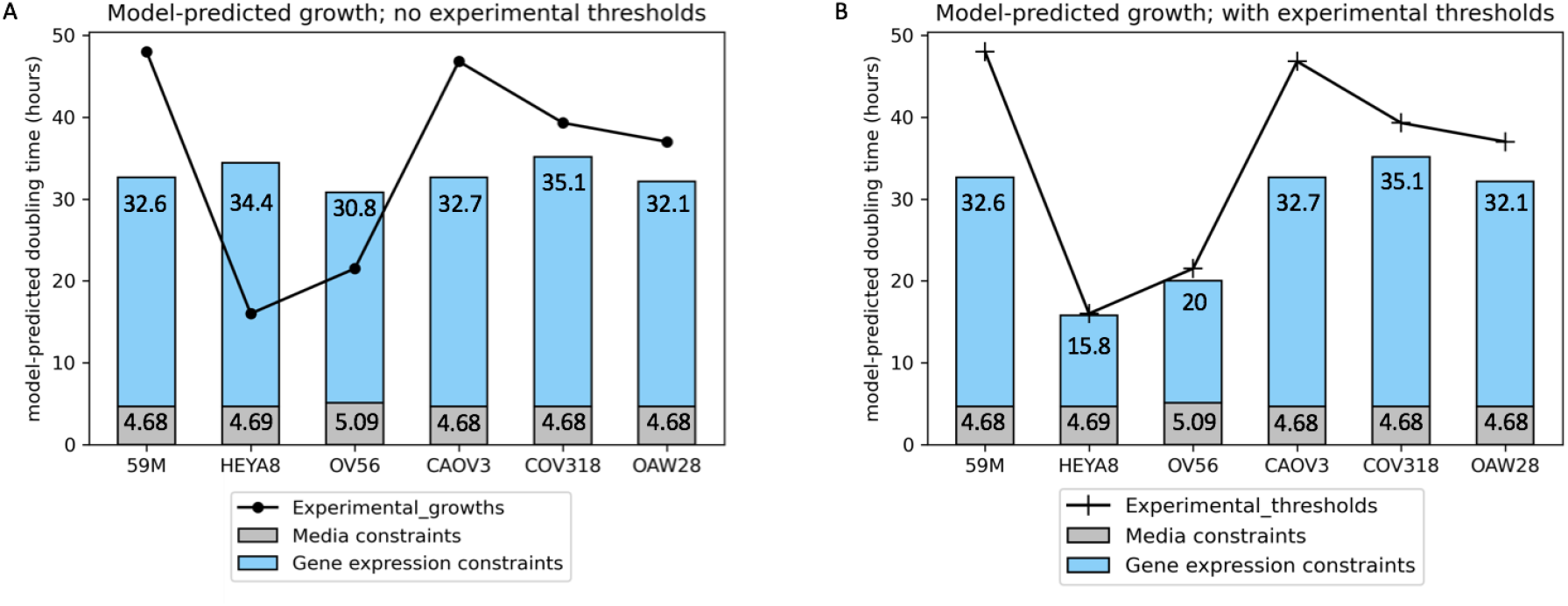
Validation of model-predicted growth rates. Experimental doubling times and their origin have been shared in Supplementary Table 2 **A.** Media constraints were applied to unconstrained Human1 using MEWpy (grey bars), and then gene expression was applied (blue bars) using our novel integration algorithm. Here, no growth thresholds were applied. **B.** Experimental growth values have been translated to thresholds for model growth predictions.

Model-predicted growths were compared, both before and after the application of experimental growths as model thresholds. Before the application of thresholds, models predicted a maximum doubling time of between approximately 30 and 35 hours, across all cell lines (Figure 3a). Experimental thresholds were most effective at tailoring model growth predictions towards experimental values in the quicker proliferating cell lines, HEYA8 and OV56 (Figure 3). On the other hand, no reactions had to be reopened for 59M, CAOV3, COV318 and OAW28, because the doubling time never reached the upper limit. In other words, gene expression did not apply tight enough constraints for the slower growing cell lines to have their growth capped, and doubling time reached a maximum across these cell lines of around 32 to 35 hours. The fact that no reactions were reopened in these four models indicated that constrained reaction bounds are entirely representative of gene expression data and media conditions. It has been concluded that using the experimental growth values as model thresholds helped generate accurate metabolic predictions. Thus, this investigation was allowed to proceed to metabolic systems flux analysis.

### Reopening reactions to satisfy experimental growth thresholds does not skew downstream metabolic predictions

Of the 13,096 reactions contained within the Human1 model, only five unique reactions had to have their constraints relaxed (bounds forced to −1000 and/or +1000 mmol/gDW/hour, depending on reversibility) across the cell lines in order to keep model-predicted doubling time below experimental maxima (Table 2). Therefore, the vast majority of fluxes analysed were representative of gene expression data. Several reactions were reopened in HEYA8 and OV56, in order to bring the model-predicted doubling time back down to the experimental maximal thresholds (Table 2). For HEYA8, there were four reactions reopened: three of these were involved in oxidative phosphorylation, whilst reaction ID MAR11443 described the transport of phenylpyruvate between the mitochondria and cytosol. Concerning OV56, there were only two reactions which were reopened – both of which were involved in oxidative phosphorylation. There are only ten reactions in the Human1 model which constitute oxidative phosphorylation, meaning that a large proportion of reaction bounds within this subsystem have been reopened to excess (−1000, 0 or 1000 mmol/gDW/hour), and in turn are not representative of cell line-specific gene expression. Therefore, conclusions made during this investigation will not involve oxidative phosphorylation. Flux predictions for these reopened reactions have been listed in Supplementary file ‘reopened_fluxes.csv’, and show that as expected, there were increased flux predictions for cell lines for which these reactions were reopened.

**Table 2.** Reactions reopened in order to keep model-predicted doubling times below experimental thresholds. For 59M, CAOV3, COV318 and OAW28 there were no reactions reopened. For HEYA8 and OV56 there were 4 and 2 reactions reopened, respectively. The majority of reopened reactions were mitochondrial and involved in the oxidative phosphorylation subsystem, as annotated by Metabolic Atlas (Robinson et al., 2020).

| Cell line | Reactions reopened | Reaction description |
| --- | --- | --- |
| 59M | N/A |  |
| HEYA8 | MAR06916 | $\text{ADP [m]} + 3 \text{ H+ [i]} + \text{Pi [m]} \Rightarrow \text{ATP [m]} + 2 \text{ H+ [m]} + \text{H}_2\text{O [m]}$ |
| | MAR06918 | $2 \text{ H+ [m]} + 2 \text{ ferricytochrome C [m]} + \text{ubiquinol [m]} \Rightarrow 4 \text{ H+ [i]} + 2 \text{ ferrocytochrome C [m]} + \text{ubiquinone [m]}$ |
| | MAR13081 | $7.92 \text{ H+ [m]} + \text{O}_2 \text{ [m]} + 4 \text{ ferrocytochrome C [m]} \Rightarrow 4 \text{ H+ [i]} + 1.96 \text{ H}_2\text{O [m]} + 0.02 \text{ O}_2^- \text{ [m]} + 4 \text{ ferricytochrome C [m]}$ |
| | MAR11443 | $\text{H+ [m]} + \text{phenylpyruvate [m]} \Leftrightarrow \text{H+ [c]} + \text{phenylpyruvate [c]}$ |
| OV56 | MAR06916 | $\text{ADP [m]} + 3 \text{ H+ [i]} + \text{Pi [m]} \Rightarrow \text{ATP [m]} + 2 \text{ H+ [m]} + \text{H}_2\text{O [m]}$ |
| | MAR06921 | $5 \text{ H+ [m]} + \text{NADH [m]} + \text{ubiquinone [m]} \Rightarrow 4 \text{ H+ [i]} + \text{NAD+ [m]} + \text{ubiquinol [m]}$ |
| CAOV3 | N/A |  |
| COV318 | N/A |  |
| OAW28 | N/A |  |

The fluxes through reactions immediately downstream of the reopened reactions shown in Table 2 have been described in Supplementary file ‘downstream_3.csv’, in order to see if the reopening of some reactions to satisfy experimentally predicted growth rates skewed overall flux results. Not all downstream fluxes were analysed, as some reopened reactions produced metabolites involved in hundreds of reactions, for example MAR06916, which produces mitochondrial ATP, is connected to 1114 reactions via its products. Results show that in general, there is no overarching trend of downstream reactions to match the flux pattern of those reopened immediately upstream. Out of the sixteen reactions identified as immediately downstream of MAR06918, which was reopened in HEYA8, there are only seven which show an increased flux in HEYA8, whilst the remaining nine reactions show either zero flux across all cell lines, or are increased in cell lines other than HEYA8. Furthermore, all three reactions immediately downstream of MAR13081 showed zero flux across all cell lines, both reactions downstream of MAR11443 have zero flux, and there was no clear pattern of upregulation in those downstream of MAR06921, which was reopened in the OV56 model. This indicates that the relaxing of a few reactions’ bounds in order to tailor growth rate towards experimental measurements does not skew the overall metabolic flux through cell line-specific models, and they are more dependent on the transcriptomics measurement – or lack of - which constrains them specifically.

### Systematic-level differences in central metabolism between low- and high-grade serous ovarian cancer

Following analysis of growth predictions, individual reaction fluxes were compared. Our significance criteria specified that there must be a 10% increase or decrease between the mean flux value for the given reaction (mean of low-versus high-grade samples), as well as at least one flux value above 0.5 mmol/gDW/hour. Of the 13,096 total reactions, 21% of the non-exchange/demand reactions had a predicted non-zero flux through at least one of the cell-lines. This meant 79% of metabolic activity was predicted to be switched off in all of the cell line-specific models. Of the non-zero subset, there were 2455 reactions which represented a 10% increase or decrease between low- and high-grade samples. The portion of this subset which had a predicted reaction flux for one or more cell lines of greater than 0.5 mmol/gDW/hour equated to 336 reactions - corresponding to 13.5% of active metabolism which showed a significant grade dependency. However, there were reactions with rates lower than this threshold of activity which represented interesting areas of metabolism. The significantly different subset has been provided in Supplementary data file ‘comparison_5.csv’.

The next step of this analysis was to gain an overview of the central metabolic systems which were enriched by these 336 significantly different reactions. Central metabolism was broadly defined according to the ‘global and overview maps’ listed in KEGG (*KEGG PATHWAY Database*, 2022), (Kanehisa & Goto, 2000), (Kanehisa et al., 2022), (Kanehisa, 2019). A database annotation search was performed in Metabolic Atlas, and of the 336 significantly different reactions, 244 were annotated with gene-reaction rules, specifying the enzyme(s) which were required for catalysis; of these there were 105 reactions belonging to central metabolic processes. Table 3 shows that nucleotide metabolism was the most enriched subsystem, with 36 reactions showing significant differences in predicted flux between low- and high-grade models. In addition, there were differences across fatty acid metabolism and the carnitine shuttle, as well as carbon metabolism, such as glycolysis, the TCA cycle, and the pentose phosphate pathway (PPP). There were significant differences predicted in DNA and RNA production, with increased flux through purine metabolism in low-grade models, and specific areas of amino acids metabolism reinforcing these changes (Table 3).

**Table 3.** Systematic overview and annotation of model-predicted significant differences. Reaction count and enzymes involved in the 105 central metabolic reactions, predicted to have significantly different fluxes (significantly up- or downregulated) between low- and high-grade models. A full list of reaction IDs has been presented in Supplementary data file ‘central_metabolism.csv’. Where the reaction may be catalysed by isoenzymes or subunit complexes, the gene encoding only one unit has been specified.

| Central metabolic subsystem | Number of reactions | Enzymes |
| --- | --- | --- |
| Nucleotide metabolism | 36 | CMPK1, GART, AK5, ADK, DCK, SLC25A19, SLC29A1, ALDH3A1, SLC25A5, NME1-NME2, RRM2B, PKM, AK2, ATIC, ADSS2, PRDX6, PAICS, PPAT, GUK1, HKDC1, PFAS, PNP, DTYMK, TXNRD2, DPYD |
| Amino acid metabolism | 23 | GOT1, GLUD1, GLUL, PRODH, PYCR3, ALDH4A1, DLD, ACADSB, ALAS1, PHGDH, VPS29, PSAT1, SHMT1, AASS, AADAT, ALDH7A1, ALDH3B1, NQO1 |
| Lipid metabolism | 13 | PNPLA4, MGLL, CDS2, GK5, GPD2, GPD1L, ALDH3A2, AKR1B1, AGK, ABHD5, SGPL1, SPHK2, PLPP1, CERS4, DEGS1 |
| Carnitine shuttle | 10 | SLC25A20, CRAT, CPT1A, SLC22A9, CROT, ELOVL5 |
| Fatty acid metabolism | 9 | SLC25A20, ACSL4, CPT1A, ALDH3A2, TECR, ACOX3, ACADVL |
| Glycolysis | 7 | PKM, DLD, GCK, LDHB, ENO1, PDHA1 |
| TCA cycle | 4 | FH, ACO2, IDH1, MDH2 |
| Pentose phosphate pathway | 3 | PGM1, PRPS2, ALDOC |

The next step was to gain understanding of the character and similarity of these overarching systems (Figure 4. Considering central carbon metabolism, glycolysis was unique in the fact that it was the only subsystem predicted to show significant upregulation in high-grade serous ovarian cancer, whilst the TCA cycle showed a 50:50 pattern of upregulation across either subtype, and the pentose phosphate pathway was entirely upregulated in low-grade models. Furthermore, there were common enzymes regulating multiple central subsystems, as represented by an overlap in circle areas in Figure 4. To be specific, enzymes regulating multiple subsystems include SLC25A20 and carnitine palmitoyltransferases, including CPT1A, which regulates the carnitine shuttle and fatty acid metabolism, as well as alcohol dehydrogenases, including ALDH3A2, which act across fatty acid metabolism and lipid metabolism. Similarly, amino acid metabolism and glycolysis share dihydrolipoamide dehydrogenase (DLD), which oxidises dihydrolipoamide to lipoamide, as well as catalysing the release of acetyl-CoA from pyruvate in glycolysis.

**Figure 4.**
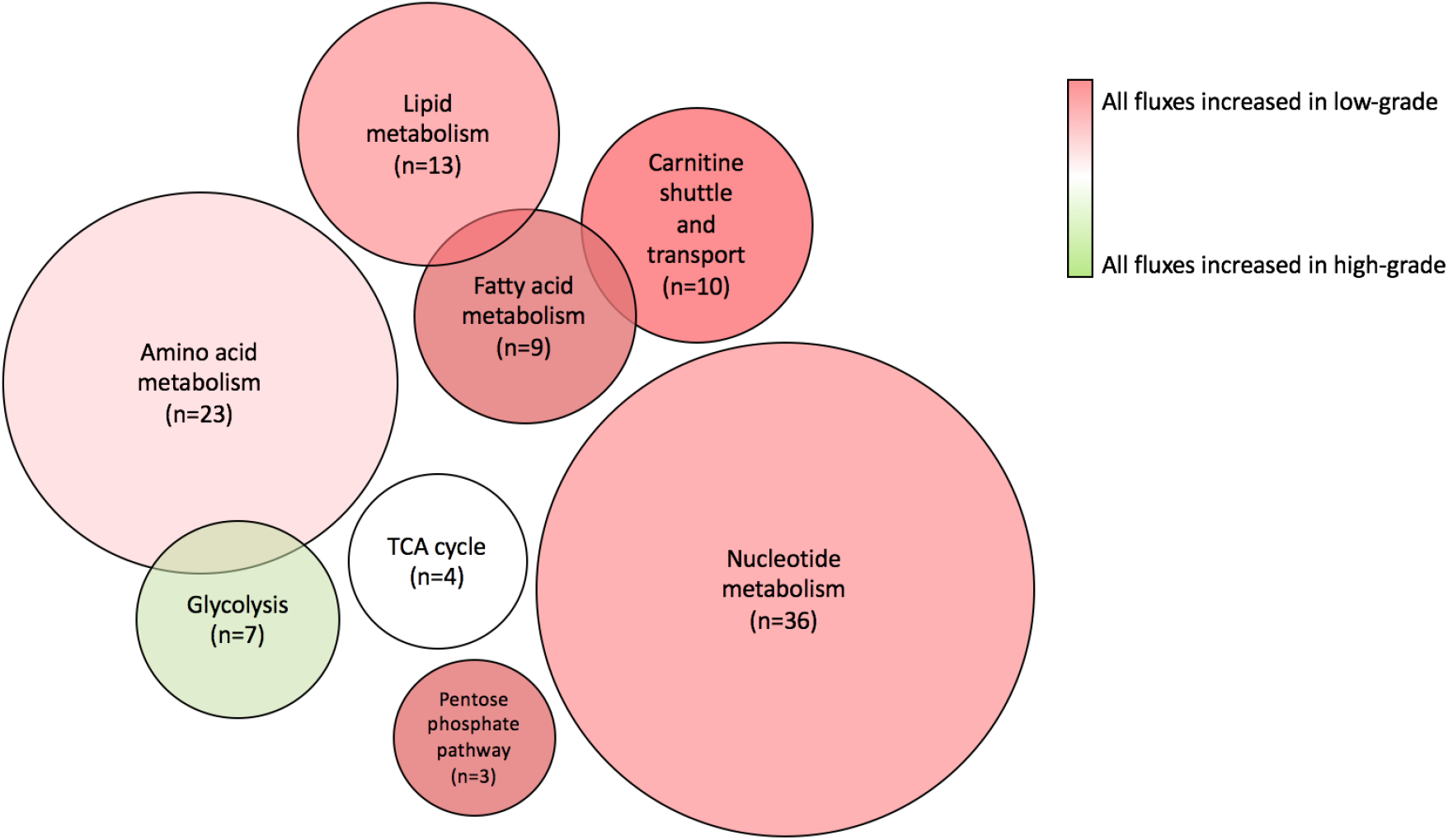
Systematic illustration of model-predicted significant differences. Venn diagram visualising the model-predicted central metabolic differences between low-and high-grade serous ovarian cancer, as listed in Table 3. Colour of circle corresponds to the subtype for which the system’s reactions have the highest flux, as indicated by key (red: upregulated in low-grade, green: upregulated in high-grade, white: 50:50 upregulation across either subtype). Area of circle corresponds to the number of significantly different reactions the model has predicted (also indicated by ‘n’ value). Overlap between circles informs that there are common enzymes in gene-reaction rules. Further details in Supplementary file ‘central_metabolism.csv’.

### Reaction-level flux differences between low- and high-grade serous ovarian cancer

After an initial, systematic view had been gained, reaction-level predictions of low- versus high-grade metabolic shifts were analysed. This analysis predicted where metabolites were accumulating, in which compartments, and, given FBA was optimised for biomass production, to what extent specific reactions could be contributing to overall cellular growth. Upon analysing the subset described above, reaction pathways were included in a final significant collection if multiple consecutive reactions were concordant in their pattern of flux, for example all reactions in a pathway were upregulated in low-grade models (Figure 5). This resulted in 41 reactions, spanning approximately 9 subsystems, as shown in Figure 5.

**Figure 5.**
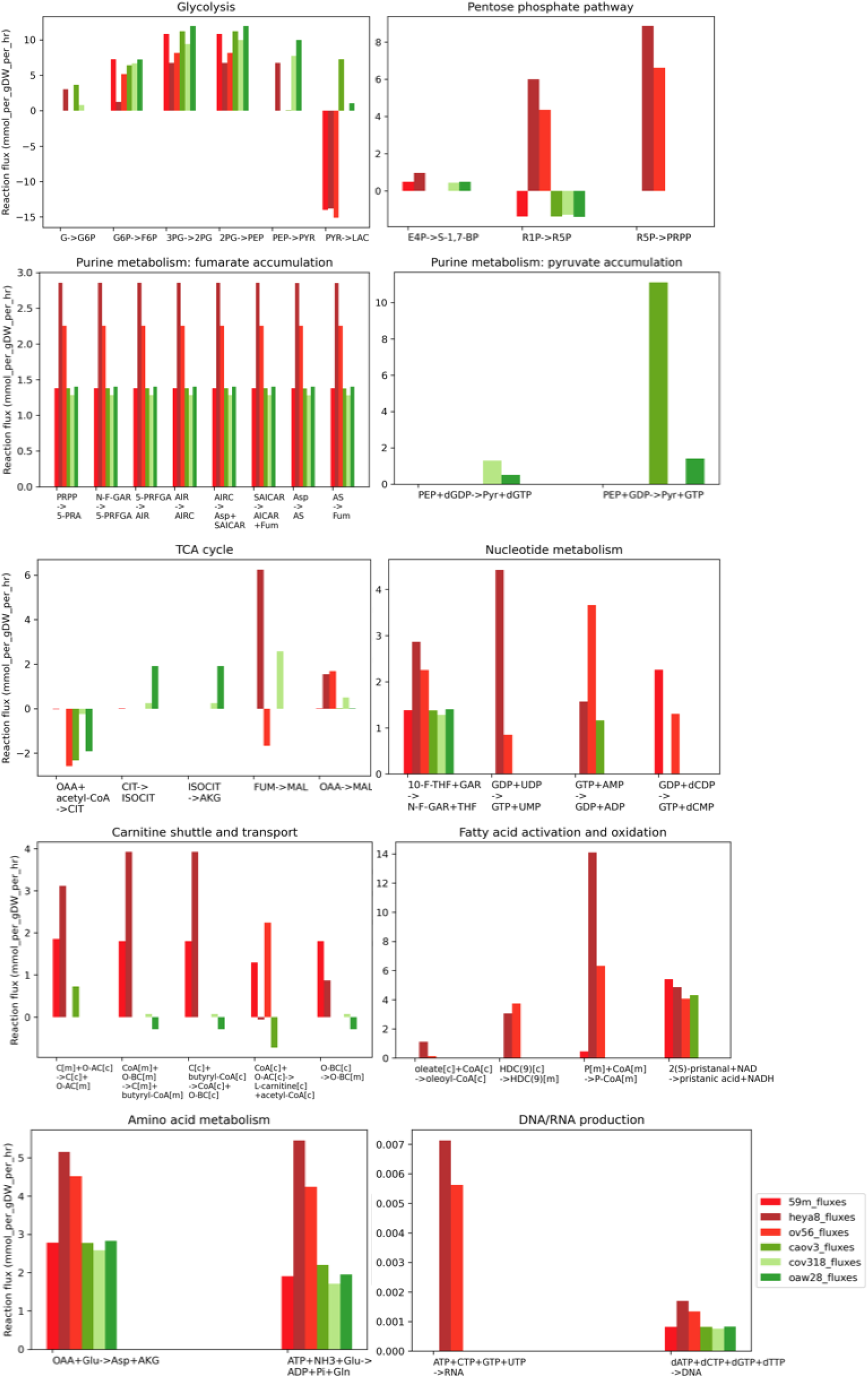
Reaction-level flux differences between low- and high-grade serous ovarian cancer. Plots to show model-predicted fluxes (mmol/gDW/hour) through central metabolism, per cell-line-specific GEM. Some of these reactions are reversible, however for clarity of visualisation, the direction shown is such that positive reaction flux corresponds to the simplified reaction equation on the x-axis. Colour of bar corresponds to cell line, as indicated in the key.

In general, Figure 5 shows a global upregulation of reaction flux in low-grade models, whilst high-grade models predicted reduced overall reaction rates. However, an exception to this rule was aerobic glycolysis, which was upregulated in the high-grade models all the way from initial glucose conversion to glucose-6-phosphate, to lactate production in the cytosol – a phenomenom which in the presence of oxygen is well-acknowledged as the Warburg effect (Warburg et al., 1927), (Vander Heiden et al., 2009).

The TCA cycle leads on from glycolysis, and models predicted differences between low- and high-grade flux, with the two-step conversion of citrate to alpha-ketoglutarate (AKG) being upregulated in high-grade, and the accumulation of malate via conversion from fumarate and oxaloacetate being upregulated in low-grade models. In order to suggest where carbon-containing metabolites may be directed in the low-grade models, if not through glycolysis, the flux through the PPP was analysed. Results suggested that fructose-6-phophate was feeding into the PPP and leading to downstream increased production of phosphoribosyl pyrophosphate (PRPP), via isomerisation of ribose-1-phosphate (Figure 5). Furthermore, differences in purine metabolism were predicted, namely increased production of fumarate in low-grade models, and increased production of pyruvate in high-grade models (Figure 5).

An increased rate of DNA and RNA production was seen in the low-grade cell lines (Figure 5). There were fluxes of 0.007 and 0.0056 mmol/gDW/hour through RNA-producing MAR07161 (Figure 5) for HEYA8 and OV56, respectively, compared to zero flux through the high-grade models. Similarly, there are fluxes of 0.0017 and 0.0013 mmol/gDW/hour through DNA-producing MAR07160 for HEYA8 and OV56, as compared to rates of less than 0.00084 mmol/gDW/hour for all high-grade models (Figure 5). Although the flux through these reactions does not exceed the initial threshold for significance (0.5 mmol/gDW/hour), these reactions have been visualised because they are downstream of significant purine metabolism reactions, and alongside other nucleotide synthesis reactions, could contribute to a cumulative significant increase in nucleic acids in the low-grade models. Finally, Figure 5 shows that for low-grade cell lines, there was increased activation of oleate to oleoyl-CoA; there was increased shuttle of O-acetylcarnitine across the IMM; increased conversion of butyryl-CoA to O-butanoylcarnitine and transport of this across the IMM, followed by conversion back to O-butyryl-CoA in the mitochondria.

### A proposed mechanism for low- versus high-grade serous ovarian cancer

The metabolic predictions mentioned thus far were amalgamated to form one mechanism depicting low- versus high-grade metabolism, and potential subtype-specific signatures. The overall mechanism includes central carbon pathways (glycolysis, PPP and TCA cycle), fatty acid metabolism (fatty acid activation, carnitine shuttle and fatty acid oxidation), amino acid metabolism, purine and nucleotide metabolism (Figure 6). The reactions predominantly occur in the cytosol, however there is also activity across the inner mitochondrial membrane, mitochondria and the nucleus.

**Figure 6.**
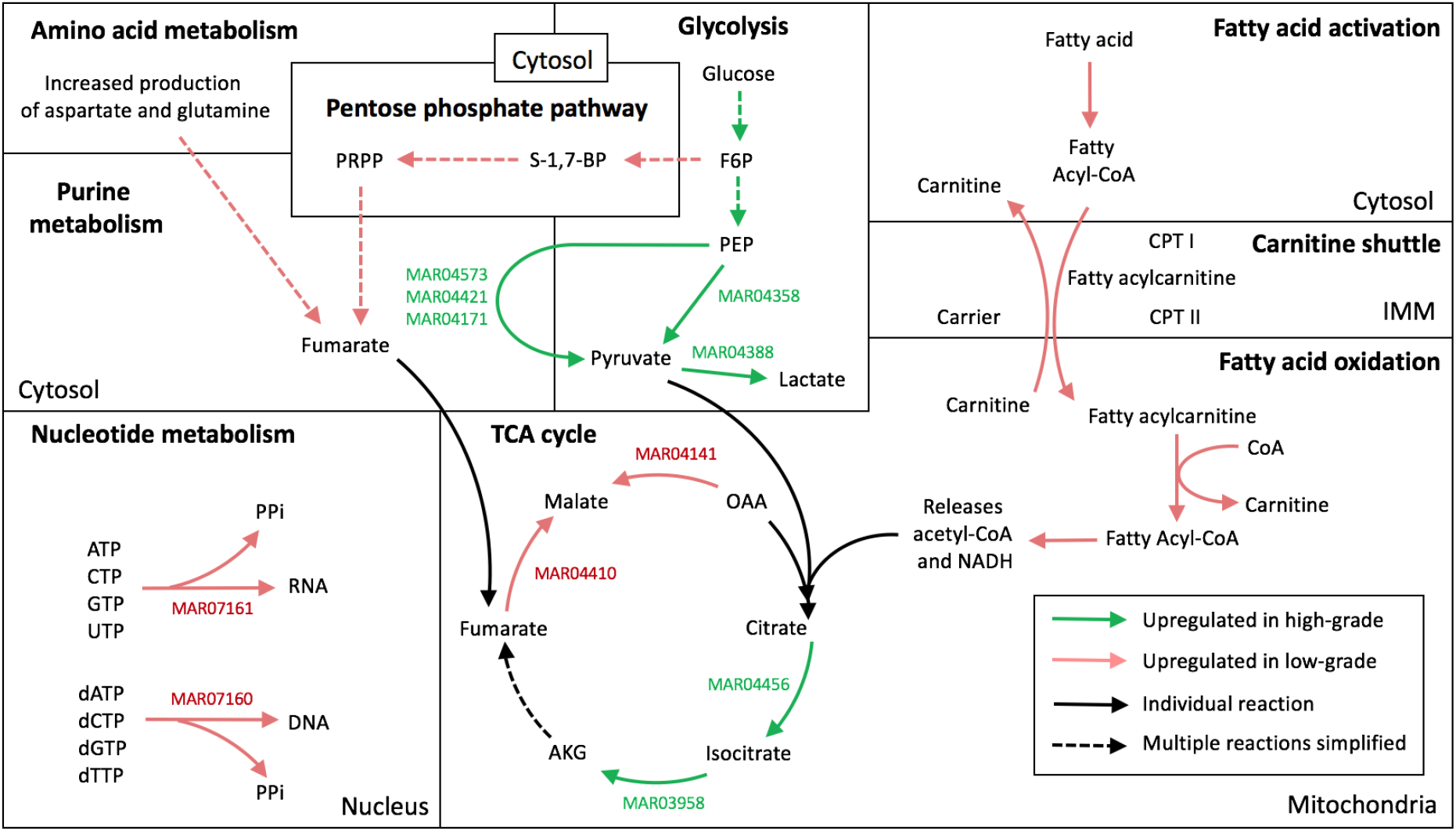
Proposed final mechanism for model-predicted low- and high-grade serous ovarian cancer differences. Compartments have been labelled, and colour of arrow denotes upregulation in LGSOC (red) and HGSOC (green). Human1 reaction IDs annotated. A solid arrow corresponds to an individual reaction, and a dotted arrow denotes multiple reactions simplified.

#### Central carbon metabolism shows differential regulation

Constraint-based modeling predicts that flux through glycolysis is increased in high-grade cellular models, fitting with the aforementioned Warburg effect – a hallmark of cancer metabolism. Pyruvate produced during aerobic glycolysis feeds into the TCA cycle via acetyl-CoA; our model suggests that pyruvate itself accumulates in high-grade models, via the purine metabolism reactions MAR04573, MAR04421 and MAR04171. In contrast, low-grade cells appear to feed fructose-6-phosphate into the PPP, with increased flux through multiple reactions, and resulting in the production of fumarate in the cytosol from PRPP – a building block for DNA and RNA. This fumarate production could be supported by increased flux through aspartate and glutamine production in the cytosol, as indicated by low-grade modeling results. Finally, multiple reactions through the TCA cycle showed differential regulation in low- versus high-grade models (Figure 6). In particular, the conversion of citrate to AKG was upregulated in high-grade models, via aconitase 1 and 2 and isocitrate dehydrogenase. However, in low-grade metabolism, models predicted an accumulation of malate from fumarate and oxaloacetate via fumarate hydratase and malate dehydrogenase 2, respectively.

#### Fatty acid metabolism and nucleic acid synthesis is upregulated in low-grade serous ovarian cancer models

According to modeling results, there is a generally increased flux through fatty acid activation in the cytosol, transport of fatty acyl-CoA molecules across the inner mitochondrial membrane (IMM) via the carnitine shuttle, and oxidation of fatty acylcarnitine to release acetyl-CoA and NADH, which can feed into the TCA cycle (Figure 6). Furthermore, there is increased flux through MAR07161 and MAR07160, which produce RNA and DNA respectively, in the low-grade models. Interestingly, the low-grade models showing the highest flux through these reactions were those with the highest experimental and model-predicted growth rates: HEYA8 and OV56. In this way, model predictions suggest that upregulation of fatty acid metabolism, purine metabolism and nucleotide metabolism could be a feature of quickly growing, low-grade cell lines.

### Gene dependency experimental data supports model predictions

In order to validate model predictions, an ORA was performed in WebGestalt, using experimental CCLE CRISPR data as an input (Ghandi et al., 2019), in order to find the predominant subsystems involving genes which show the most contrasting dependency scores between low- and high-grade ovarian cell lines. Of the mean gene dependency scores (low-grade average, and high-grade average), the 200 gene IDs which showed the highest dependency in low-grade cell lines were used as input for the ORA. Of these 200 IDs, 196 could be mapped to Entrez gene IDs, and 91 of these IDs could be annotated with functional categories in WebGestalt. The most enriched pathway was fatty acid biosynthesis, with three genes identified from the mapped input: ACACA, ACSL3 and OXSM, and an enrichment ratio of 18.94 (Figure 7a). The second most enriched pathway was the pentose phosphate pathway, with five genes identified from the mapped input (Figure 7***b***).

**Figure 7.**
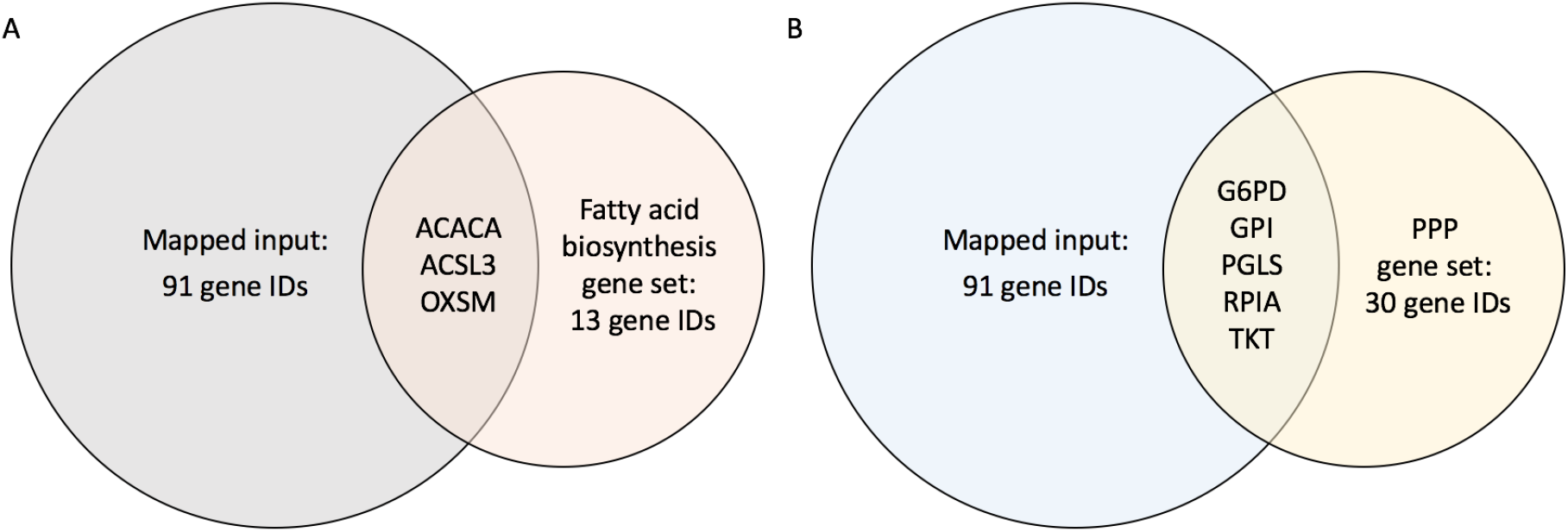
Over-representation analysis for low-grade specific growth pathways. ORA was performed in WebGestalt (Liao et al., 2019), to find the over-enriched pathways for genes showing the highest dependency scores in low-grade cell lines. Gene dependency scores were averaged for low-grade (59m and HEYA8) and high-grade (CAOV3, COV318 and OAW28) cell lines, and the top 200 showing the highest value in low- compared to high-grade cell lines was used as input for ORA. Enrichment category was pathway and KEGG databases, with genome platform as the chosen gene reference list. Number of genes successfully mapped in input and contained in gene sets are annotated on bubble. Overlapping genes highlighted in center of bubbles **A.** The most over-enriched pathway was fatty acid biosynthesis, with three genes found in the mapped input. False discovery rate (FDR) was 0.029867, p-value was 4.5 × 10^−4^ and enrichment ratio was 18.94. (ACACA; acetyl-CoA carboxylase alpha, ACSL3; acyl-CoA synthetase long chain family member 3, OXSM; 3-oxoacyl-ACP synthase, mitochondrial). **B.** The second most over-enriched pathway was pentose phosphate pathway (PPP), with five genes from the mapped input. FDR was 0.0087792, p-value was 2.7 × 10^−5^ and enrichment ratio was 13.68. (G6PD; glucose-6-phosphate dehydrogenase, GPI; glucose-6-phosphate isomerase, PGLS; 6-phosphogluconolactonase, RPIA; ribose 5-phosphate isomerase A, TKT; transketolase).

Assuming that genes with high dependency scores upregulate their corresponding model reactions to a greater extent, the CCLE gene dependency data agrees with model predictions. In the cell-line specific models, ACACA catalyses fatty acid biosynthesis (MAR04156) and biotin metabolism reactions (MAR07672 and MAR07673), all of which have a higher average flux value in low- compared to high-grade cell lines. Furthermore, ACSL3 is involved in fatty acid activation reactions, which have been shown to have higher fluxes in the low-grade specific models (see Figure 6). Similarly, OXSM facilitates the co-transport of butyryl-CoA and carnitine between the mitochondria and the cytosol (MAR00158), and this reaction is upregulated in low-grade-specific models (average fluxes of 1.9 mmol/gDW/hour versus 0.024 mmol/gDW/hour) (see Figure 6). the second most enriched pathway for the genes with highest dependency scores in low-grade cell lines was the pentose phosphate pathway, which is also predicted to be upregulated in a low-grade-specific manner (see Figure 5 and Figure 6), further validating model predictions. Of the five enzymes pulled from the mapped input, G6PD, PGLS, RPIA and TKT showed general patterns of increased flux through low-grade-specific models, highlighting why the PPP was suggested to be a preferred metabolic avenue for growth in low-grade cell lines (see Figure 6).

As well as the ORA, we performed an *in silico* KO simulation to see if models could replicate the growth patterns reported in the CCLE CRISPR growth dependency dataset. Here, single genes were knocked out, and growth was calculated before and after, to show which genes were essential for cell lines to reach optimal growth rates. Experimental growth dependency scores were correlated with these ratios, and of the five cell lines which were available in the CCLE gene dependency dataset (59M, HEYA8, CAOV3, COV318 and OAW28), there was a general pattern of negative correlation (values between −0.319 and −0.356), showing that as experimental growth dependency scores increased, model simulations predicted that knockout of the same gene would be detrimental for growth. Given the complexity and scale of human genome-scale models, it can be difficult to demonstrate sensitivity of cellular growth to genetic changes, therefore, this correlation analysis was extended to focus on predicted essential genes. For the cell lines for which *in vitro* gene dependency data was available, genes which had no predicted essentiality (as indicated by a before/after growth ratio of 1.0) were removed from the initial knockout simulation dataset. From here, there were between 267 and 305 essential genes predicted per cell line, and correlation of KO simulation data with experimental data became more negative, between −0.326 and −0.458, with an average increase (towards a more negative value) of 20% per cell line. Considered together, these validations show the models are able to predict features of metabolism which support the growth of ovarian cancer, in a subtype-specific manner, and KO simulations show agreement with experimental gene dependency data, for predicted essential reactions.

## Discussion

Results presented here indicate a clear difference in the metabolism of low- and high-grade serous ovarian cancer, with our analysis predicting that of the active metabolism (roughly 20% of total reactions), 13.5% of reactions shows a significant subtype-dependency. Of this significant subset, 43% represented central metabolism. When this overview was divided into individual subsystems, many of the model’s predictions were concordant with that which has been reported for cancer metabolism in general, and also for ovarian subtype-dependent signatures. Furthermore, metabolic predictions were able to explain clinical phenotypes observed, for example metastasis and chemoresistance.

### Glycolytic upregulation correlates with aggressive phenotype

Cell line-specific models predicted that aerobic glycolysis is upregulated in HGSOC models, extending all the way to increased lactate production. Similar modeling studies have been performed linking metabolism to clinical phenotype. For example, one genome-scale investigation compared early-stage and more aggressive, late-stage breast cancer, and associated increased glycolysis with the more aggressive phenotype (Jerby et al., 2012). In addition, it was suggested that there exists a ‘go or grow’ dichotomy in breast cancer, where cells exhibit a trade-off between proliferation rates, which are increased in early-stage, and increased metastatic capability, which is greater in late-stage breast cancer (Jerby et al., 2012). This connection raises the question, could a similar phenomenon be present in ovarian cell lines, where increased growth rates did in fact correlate with a drop in glycolytic rate?

This association between glycolysis and an aggressive phenotype has also been reviewed in the context of ovarian cancer (Nantasupha et al., 2021). Increased glycolysis in ovarian cancer is described to support cellular migration and survival through increased formation of F-actin via phosphoglycerate mutase 1, and regulation of mTOR and apoptosis via hexokinase 2 and glyceraldehyde-3-phosphate dehydrogenase (Nantasupha et al., 2021). In this way, the observation of increased glycolysis in high-grade samples reported here is supported by literature, and could contribute to the mechanistic understanding of tumour behaviour.

### Deregulated fatty acid metabolism supports low-grade proliferation and chemoresistance

On top of perturbations to glycolysis, fatty acid metabolism and its biosynthesis in particular were pathways which models predicted to be upregulated in low-grade cell lines. As described, this has been validated using experimental gene dependency data and knockout simulations. The way in which fatty acid oxidation is connected to carbon metabolism is through the generation of reducing equivalents, such as NADH and acetyl-CoA, which feed into the TCA cycle. In fact, the same investigation which linked increased glycolysis to late-stage breast cancer, described increased fatty acid biosynthesis as a feature of low-grade tumours, which supports their increased proliferation (Jerby et al., 2012). Therefore, it could be suggested that low- and highgrade ovarian tumours behave metabolically in a manner parallel to early- and late-stage breast cancer, respectively. It has been suggested that upregulation of lipid metabolic enzymes is a metabolic hallmark of cancer cells (Koundouros & Poulogiannis, 2020), but here it has been shown that genome-scale modeling bridges the gap between these observed gene expression signatures and actual metabolic significance. Evidence from prostate cancer studies suggest a parallel phenotype to that which has been predicted here, and explains that clinically, glycolysis is downregulated in prostate cancers, and this is partnered with the idea that β-oxidation is the main bioenergetic pathway for these cells (Koundouros & Poulogiannis, 2020). This comparison serves to validate and explain the contrasting glycolytic and lipid regulation predicted in these low-grade ovarian cancer models.

It is known that fatty acid metabolism supports cellular proliferation and tumorigenesis, however it remains unknown how this relates to chemoresistance – one of the major challenges in low-grade serous ovarian cancer. Multiple associations have been made between lipid metabolism and chemoresistance in ovarian cancer, for example, FABP4 expression has been shown to increase lipolysis and lead to carboplatin resistance (Germain et al., 2020). Similarly, increased fatty acid synthase (FASN) expression promotes lipogenesis and causes cisplatin, carboplatin and paclitaxel resistance in ovarian cancer (Germain et al., 2020), (Yang et al., 2022).

### Possible co-regulation of the pentose phosphate pathway and nucleotide synthesis in low-grade models

Modeling predictions suggest that the pentose phosphate pathway is a favourable avenue for growth in low-grade models and this has been validated with functional annotation analysis on experimental gene dependency data. In addition to the regulation of glycolysis and fatty acid metabolism which appear to overlap between breast cancer and ovarian predictions, nucleotide biosynthesis shows similar patterns of flux. It was reported that nucleotide biosynthesis is upregulated in early-stage breast cancer to support proliferation (Jerby et al., 2012), and this parallels the low-grade ovarian models presented here. The final mechanism illustrated in Figure 6 suggests that an increase in the pentose phosphate pathway – defined by its role in producing precursors for nucleotide synthesis – could support nucleotide biosynthesis in low-grade cell lines. Furthermore, modeling results suggest that these low-grade observations are accompanied by an increase in purine synthesis, as supported by aspartate and glutamine production. Nucleotides may be synthesised via the salvage or de novo synthesis pathways. Salvage includes the recycling of nucleosides from DNA and RNA degradation, and de novo synthesis includes simple sugars and amino acids as building blocks. In general, quickly dividing cancer cells rely more on de novo nucleotide synthesis (Liu et al., 2022), which, alongside the salvage pathway, is dependent on PRPP (a product from the PPP). This could explain why the PPP is predicted to be upregulated alongside nucleotide metabolism for low-grade, quickly dividing ovarian cell lines. Furthermore, adenylate kinases 4 and 7 (AK4/7), which balance cellular ATP levels, are of particular focus for ovarian research (Liu et al., 2022). Model predictions show a significant increase in activity through all three reactions catalysed by AK4/7 (MAR04002, MAR04004 and MAR04480) in low-grade cell lines, and research has associated AK4-enhanced hypoxia with chemoresistance (Liu et al., 2022), a feature of low-grade ovarian tumours.

### Targeting the metabolic signatures of low- and high-grade serous ovarian cancer

In addition to predicting metabolic signatures, one purpose of genome-scale modeling is to provide the framework or hypotheses for future drug discovery research. A perfect example of this is a genome-scale modeling investigation which employed HMR2 in the context of hepatocellular carcinoma, and ran constraint-based simulations to predict antimetabolites *in silico*, and then impressively, found that around one fifth of the predicted antimetabolites were already in use in cancer treatment strategies (Agren et al., 2014).

There exists an abundance of evidence regarding the targeting of ovarian tumour metabolism. For instance, a prediction of high-grade metabolism was increased glycolytic rates, and studies have demonstrated that owing to the increased expression of glycolytic enzymes in HGSOC (GLUT1 and HKII), cell proliferation may be reduced using a panel of glycolytic pathway inhibitors (Xintaropoulou et al., 2018). Furthermore, the same study showed that this inhibition was effective in attenuating growth in both platinum-sensitive and resistant cell lines, and novel GLUT1 and LDH inhibitors demonstrated synergy with metformin – an inhibitor of oxidative phosphorylation (Xintaropoulou et al., 2018). Finally, the cell line panel from which these *in vitro* results were generated contained CAOV3 – one of the high-grade cell lines modelled in this project, thus suggesting *in silico* glycolytic engineering experiments could yield promising results for CAOV3.

A predicted metabolic signature of low-grade cell lines was the upregulation of the PPP, to feed into nucleotide biosynthesis. This feature could represent a vulnerability in low-grade, chemoresistant cell lines. A study showed that some cancer cells rely on the pentose phosphate pathway, to synthesis nucleotides and support their proliferation, and that inhibition of the PPP (via 6-aminonicotinamide and dichloroacetate) was able to limit cellular proliferation (De Preter et al., 2015). Furthermore, modeling results have predicted the upregulation of fatty acid metabolism in low-grade cell lines, and research has listed many examples of how we could exploit vulnerabilities in lipid metabolism, and potentially target chemoresistance. For example, there are preclinical drugs (G28UCM and Orlistat) which target FASN, and act by increasing cytotoxicity and suppression of HER2 overexpression (Germain et al., 2020). In addition, an inhibitor of FABP4 has been described (BMS309403), which reduces tumour burden and increases the sensitivity towards carboplatin in cell lines and *in vivo* models (Germain et al., 2020).

### Important *in silico* developments

Despite the translational potential of cell line-specific models there remains some limitations of genome-scale modeling that may be overcome with future *in silico* developments. For instance, FBA requires an objective function to be defined, meaning all flow through reactions in the model is optimised for one cellular task. Biomass production, in general, is a good objective function for modeling cancer cells, due to replicative immortality being a hallmark of cancer. The definition of the objective function affects gene essentiality analysis, which usually categorises genes as essential if, upon their deletion, biomass production is reduced to zero. However, as we know, human cells are specialised for a range of metabolic tasks, as shown by the discussion into the definition of gene essentiality accompanying the release of the Human1 model (Robinson et al., 2020). Here, as opposed to solely impairing cellular growth, a gene was defined as essential if it impaired any of 57 basic metabolic tasks, including functions such as de novo synthesis of ATP, oxidative phosphorylation and β-oxidation (Robinson et al., 2020).

A potential future update to the methods described here would be the incorporation of proteomics data, to generate a multi-omics model of ovarian cancer metabolism. There is poor correlation between gene and protein expression, and transcriptomics has been proven to harbor limited predictive power for actual protein abundance (Gygi et al., 1999), (Nusinow et al., 2020). Despite the poor correlation, transcriptomics has been used for this project due to its high-throughput nature (Z. Wang et al., 2009), which may overshadow its limitations in discounting the impact of post-translational modifications, protein degradation and other factors which give rise to contrasting gene-to-protein measurements. We have compensated for this by relaxing our reaction bounds to satisfy experimental growth rates, to deduce areas of gene expression which may not accurately reflect the proteome and growth phenotype. Furthermore, a multi-omics approach in ovarian research has permitted an integrative analysis of signaling pathways, and aided precision medicine spanning from genetic drivers to their downstream protein effectors (Khella et al., 2021).

Finally, initial analyses reported here were limited to only six cell line-specific models, because the defined media was standardised in order to ensure the models were reflecting gene expression (rather than media-related) metabolic differences. Furthermore, the composition of FBS could only be estimated, using the reaction closing and reopening which has been described, due to a lack of published definition of this media. Therefore, future models would have their accuracy and reliability improved by using a better defined media, or by incorporating a full chemical characterisation of FBS.

### Concluding remarks

Here, we have developed an ‘omics integration method for constraint-based modeling, using the Human1 GEM framework and ovarian cancer gene expression data. This work has shown that the use of experimental growth thresholds, and the relaxing of reaction bounds, can compensate for inconsistencies between gene expression and growth phenotype – a method which has shown high efficacy in quickly proliferating cell lines. Furthermore, metabolic predictions, namely the upregulation of glycolysis in high-grade, and increased flux through the PPP, nucleotide metabolism and fatty acid metabolism in low-grade serous ovarian cancer, are concordant with literature evidence. On top of this, experimental growth dependency scores have been able to validate the predicted low-grade-specific essentiality of the pentose phosphate pathway and fatty acid biosynthesis, with KO simulations demonstrating the accuracy of constraint-based modeling to predict gene essentiality. The potential metabolic vulnerabilities of either subtype have already been targeted with preclinical drugs, indicating they may represent legitimate ovarian cancer signatures. In addition, results generated here could contribute to a greater understanding of the metabolic heterogeneity of ovarian cancer, and the specific features responsible for contrasting chemosensitivities and survival prognosis observed for low- and high-grade subtypes. This method could be translated to other diseases and datasets, and these cell line-specific models could provide a framework for future genetic engineering experiments and drug simulations, inspiring personalised medicine studies and improving outlook for ovarian cancer patients.

## Supporting information

central_metabolism

comparison_5

downstream_3

lg_hg_media_subtype_site

reopened_fluxes

unannotated

