## Supplementary figures for "Constraint-based modeling predicts metabolic signatures of low- and high-grade serous ovarian cancer"

### Supplementary data

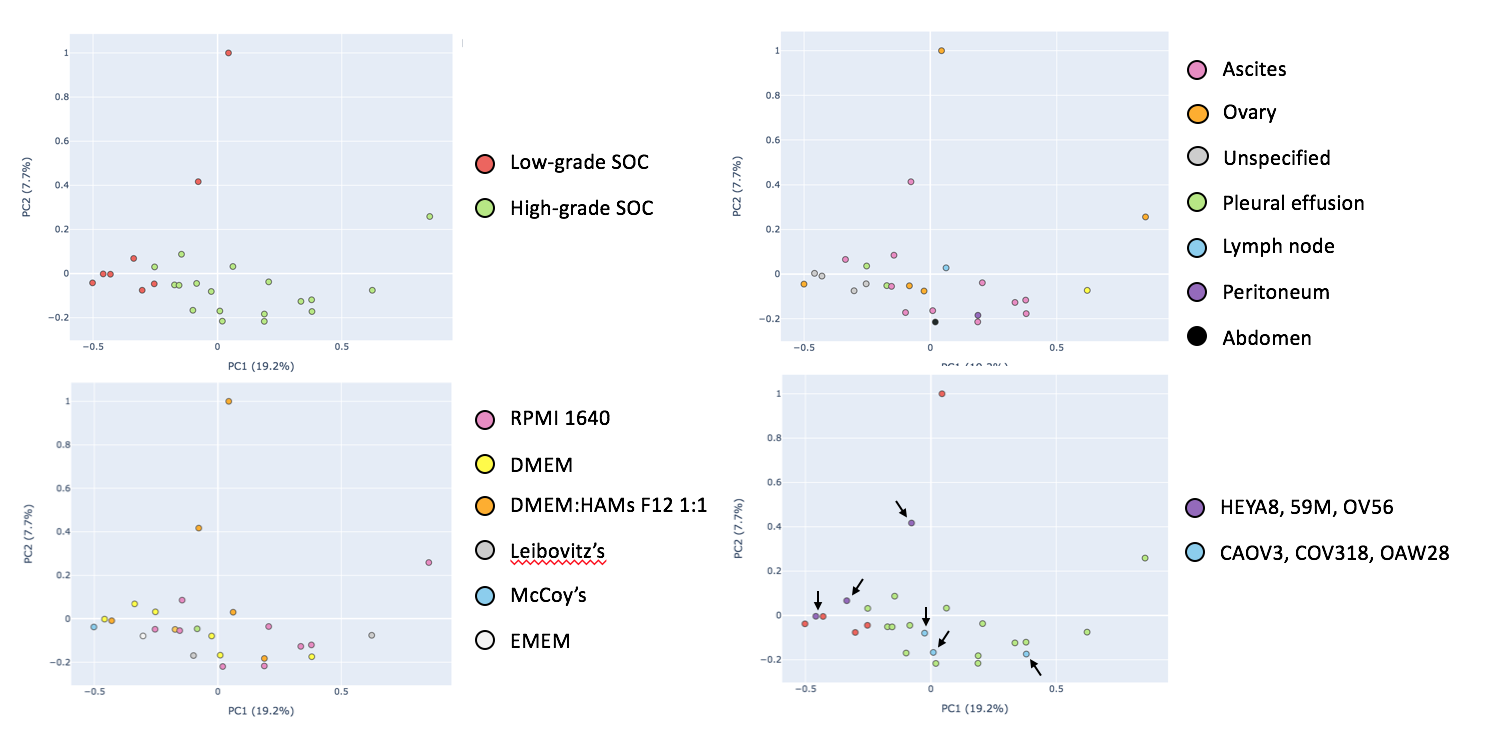

**Supplementary Figure 1. Visualising how media and site of origin control RNAseq pattern.**

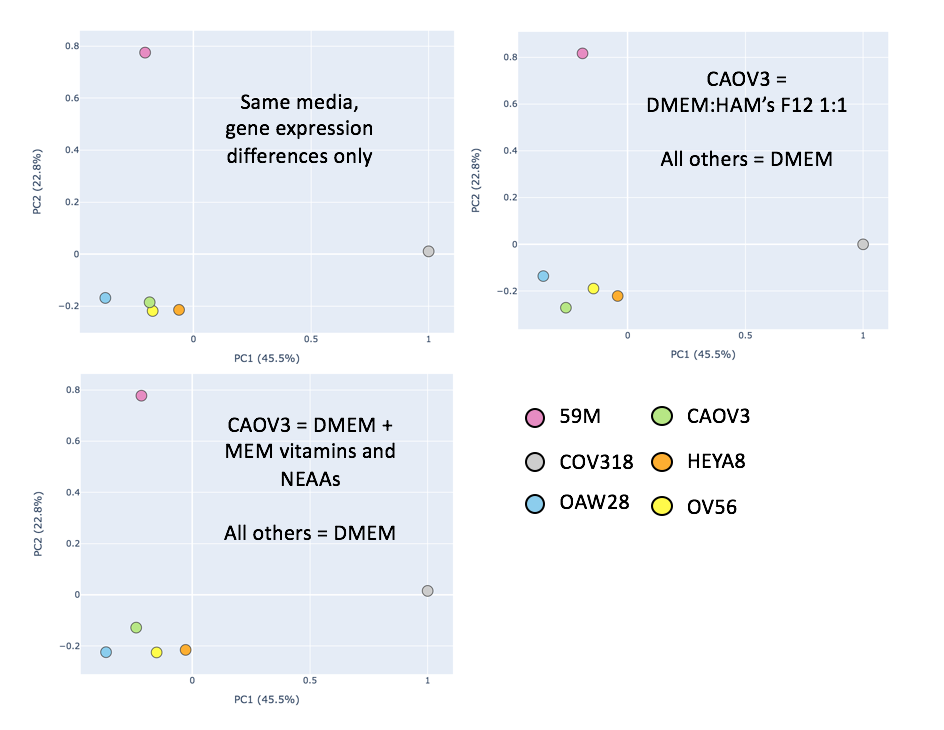

**Supplementary Figure 2. Standardisation of media conditions.**

**Table 1. Media conditions for chosen CCLE cell lines.**

| Cell line | Media composition | Reference for media composition |
| --- | --- | --- |
| 59M | DMEM + 2mM Glutamine + 1mM Sodium Pyruvate (NaP) + 20 IU/l Bovine Insulin +10% FBS | (Wilson, 2022a) |
| HEYA8 | DMEM + 1% MEM vitamins + 1% MEM non-essential amino acids +10% FBS + Penicillin-Streptomycin | (Haley et al., 2016) |
| OV56 | DMEM:HAM’s F12 1:1 +5% FBS 0.5 ug/ml hydrocortisone + 10ug/ml insulin | (Barretina et al., 2012) |
| CAOV3 | DMEM +10% FBS https://www.atcc.org/products/htb-75 | (Tsherniak et al., 2017) |
| COV318 | DMEM +10% FBS +2mM glutamine | (Barretina et al., 2012) |
| OAW28 | DMEM + 2mM Glutamine + 1mM Sodium Pyruvate (NaP) + 20 IU/l Bovine Insulin +10% FBS | (Wilson, 2022b) |

**Table 2. Experimental and model-predicted growth rates.**

| Cell line | Experimentally-predicted growth rate | Source of experimental growth rate | Model-predicted growth rate |
| --- | --- | --- | --- |
| 59M | 48 | Cellosaurus.org | 32.62 |
| HEYA8 | 16 | Cellosaurus.org | 15.77 |
| OV56 | 21.5 | Taylor lab (personal communications) | 20 |
| CAOV3 | 46.8 | Taylor lab (personal communications) | 32.67 |
| COV318 | 39.3 | Taylor lab (personal communications) | 35.13 |
| OAW28 | 37 | Cellosaurus.org | 32.12 |

### Supplementary files

comparison_5.csv

central_metabolism.csv

unannotated.csv

reopened_fluxes.csv

downstream_3.csv

lg_hg_media_subtype_site.csv
